# Auditory cortex is susceptible to lexical influence as revealed by informational vs. energetic masking of speech categorization

**DOI:** 10.1101/2020.10.20.347724

**Authors:** Jared A. Carter, Gavin M. Bidelman

## Abstract

Speech perception requires the grouping of acoustic information into meaningful phonetic units via the process of categorical perception (CP). Environmental masking influences speech perception and CP. However, it remains unclear at which stage of processing (encoding, decision, or both) masking affects listeners’ categorization of speech signals. The purpose of this study was to determine whether linguistic interference influences the early acoustic-phonetic conversion process inherent to CP. To this end, we measured source level, event related brain potentials (ERPs) from auditory cortex (AC) and inferior frontal gyrus (IFG) as listeners rapidly categorized speech sounds along a /da/ to /ga/ continuum presented in three listening conditions: quiet, and in the presence of forward (informational masker) and time-reversed (energetic masker) 2-talker babble noise. Maskers were matched in overall SNR and spectral content and thus varied only in their degree of linguistic interference (i.e., informational masking). We hypothesized a differential effect of informational versus energetic masking on behavioral and neural categorization responses, where we predicted increased activation of frontal regions when disambiguating speech from noise, especially during lexical-informational maskers. We found (1) informational masking weakens behavioral speech phoneme identification above and beyond energetic masking; (2) low-level AC activity not only codes speech categories but is susceptible to higher-order lexical interference; (3) identifying speech amidst noise recruits a cross hemispheric circuit (AC_left_ → IFG_right_) whose engagement varies according to task difficulty. These findings provide corroborating evidence for top-down influences on the early acoustic-phonetic analysis of speech through a coordinated interplay between frontotemporal brain areas.

## 1. INTRODUCTION

Prior to engaging in the cognitive processes of speech communication, an individual must map sensory cues onto a perceptual map through the process of categorical perception (CP). In terms of speech perception, acoustic signals vary continuously across a wide range of spectral features, but the transformation of acoustics properties to discrete phonetic labels allows listeners to use sounds in their auditory-linguistic system to engage in communication (Liberman et al., 1967; Pisoni, 1973). CP is demonstrated when listeners perceive gradually morphed speech signals along a continuum are heard as one of only a few discrete phonemes, a process of perceptual “downsampling” or grouping.

Previous neuroimaging work using event related potential (ERPs) has shown neural correlates of categorization when comparing response amplitudes to prototypical vs. category-ambiguous speech tokens, with effects occurring as early as 150-200 ms after stimulus presentation (Bidelman et al., 2013; Bidelman and Walker, 2017; Liebenthal et al., 2010; Mankel et al., 2020). The introduction of acoustic degradation (i.e., noise) prolongs and weakens the ERPs, indicating predictable masking on speech processing (Bidelman and Howell, 2016; Bidelman et al., 2018; Billings et al., 2009). However, we have recently shown that speech categories—those carrying a strong phonetic identity—are more resilient to noise degradation than their phonetically ambiguous counterparts (Bidelman et al., 2020a). Categorization recruits a wide variety of frontal, temporal, and parietal brain regions (Al-Fahad et al., 2020; Chang et al., 2010; Myers et al., 2009). Yet, category-level representations are highly prominent in inferior frontal gyrus (IFG) (Bidelman and Walker, 2019; Myers et al., 2009) and auditory cortex (AC) (Bidelman and Lee, 2015; Bidelman and Walker, 2019; Chang et al., 2010), suggesting fronto-temporal interplay is an important driver of sound labeling. In the context of degraded speech, we also know that the relative involvement of IFG and AC to speech coding varies with the severity of noise interference (Bidelman and Howell, 2016; Du et al., 2014). Moreover, when masking is introduced in speech tasks, frontal lobe activates bilaterally (Price et al., 2019; Scott et al., 2004), suggesting an integrated network in which frontal regions—particularly IFG for phonetic tasks—helps resolve ambiguity in the signal caused by the masker.

One debated issue in speech perception and CP is how higher-order linguistic representations inform sub-lexical processing (i.e., phonemes). One end of this spectrum suggests categorization is hard-wired into neurophysiology, evidenced by animals with no linguistic capability successfully categorizing speech phonemes (Kuhl, 1986). This model suggests rigid prototypes of speech that other factors surrounding the phoneme (e.g., context, listener expectations, later phonemes, etc.) cannot influence perception. However, evidence exists that top-down linguistic processing affects speech categorization, causing category representations to be malleable. Indeed, speech categories change with variations in context (e.g., Ganong effect), with language or music experience, and from non-linear dynamics in perception (Bidelman et al., 2020b; Ganong, 1980; Mankel et al., 2020; Tuller et al., 1994; Tuller, 2005). Thus, while bottom-up encoding initiates CP, top-down effects reveal that the early formation and perceptual representations for speech phoneme categories are shaped by a complex auditory-linguistic network.

Other converging evidence for top-down influences in CP are effects of informational masking on spoken word comprehension. While energetic masking (i.e., peripheral masking based on cochlear mechanics) can degrade the speech signal, informational masking describes perceptual interference due to a similarity or confusability of target and masker sounds—a central-cognitive aspect of figure-ground perception (Kidd et al., 2008). Indeed, parsing target speech from concurrent speech (a form of informational masking) is more challenging to comprehension than when maskers are non-linguistic in nature (e.g., broadband noise), even after controlling for spectral and/or SNR differences between the maskers (Krizman et al., 2017; Swaminathan et al., 2015; Yoo and Bidelman, 2019). As this effect cannot be attributed to bottom-up encoding of the signal, informational masking necessarily reflects top-down influences in speech perception. Within the brain, informational masking has been shown to influence the auditory ERPs and neural differentiation of speech sounds above and beyond energetic masking alone (Bennett et al., 2012; Carter, 2018; Niemczak and Vander Werff, 2019). These findings suggest higher-order linguistic brain areas likely influence early perceptual phoneme coding in lower-order (canonical) auditory brain areas, particularly when informational masking stresses the system. Under investigation here is when and where these top-down influences on speech perception occur in the context of categorization and the acoustic-phonetic conversion process inherent to CP. We were also interested to test whether energetic vs. informational masking exerts a differential effect on category formation (cf. Bidelman et al., 2020a), such that speech categories form in different brain regions (i.e., AC vs. IFG) depending on concurrent lexical interference.

To this end, the current study aimed to: (1) evaluate whether linguistic interference (measured via informational masking) influences the early acoustic-perceptual conversion inherent in CP and if so, what stage of processing (e.g., AC vs. IFG or encoding or decision, respectively) they occur; (2) identify the neural circuitry used to resolve ambiguity in multiple streams of competing speech. To address these questions, we measured behavior, source-level ERPs, and functional connectivity in young adults during rapid phoneme identification tasks using a five-step consonant-vowel (CV) continuum (**Figure 1**) in the presence of different types of noise (energetic vs. informational masks). If top-down lexical processing impacts phonetic coding, we hypothesized a differential (stronger) effect of informational masking on categorization compared to matched energetic masking with similar spectral characteristics, with perhaps increased activation of frontal regions when disambiguating the signal from the masker. Our findings reveal auditory cortex not only codes speech categories but is susceptible to higher-order lexical interference under the stressors of informational masking.

**Figure 1:**
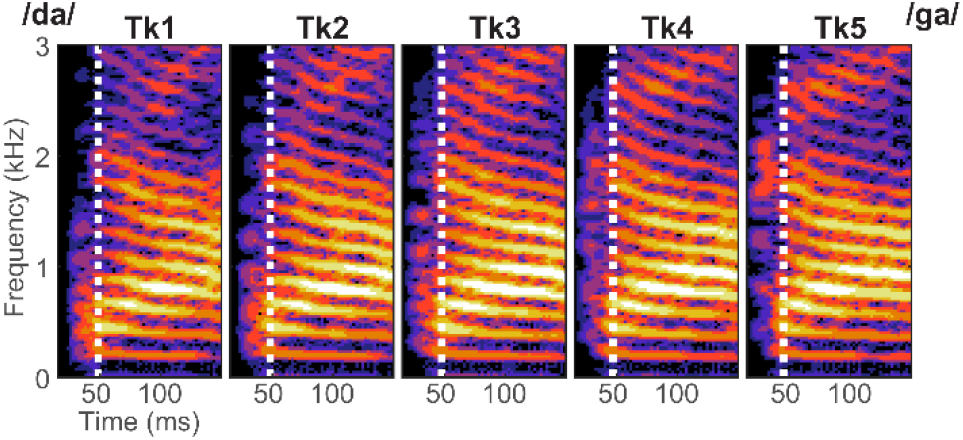
Acoustic spectrograms for the /da/ to /ga/ speech continuum. Tokens were spaced over five equidistant steps by varying the F2 formant using STRAIGHT (Kawahara et al., 2008) based on original speech materials described in Nath and Beauchamp (2012). Effective token duration was 350 ms (zoomed here for clarity). Speech was presented at 74 dB SPL.

## 2. RESULTS

### 2.1 Behavioral data

Behavioral identification functions are shown for the different masking conditions in **Figure 2A**. Listeners perceived the tokens categorically in the clean condition, but the addition of noise caused categorization to become more continuous, as evidenced by the flattening of the psychometric functions. Analysis of the slopes showed a main effect of masking (*F*_2,28_ = 46.89, *p* < 0.0001) (**Figure 2D**). Tukey-Kramer contrasts revealed that relative to clean speech, the addition of both informational (*p* < 0.0001) and the energetic (*p* < 0.0001) masking hindered categorization. More critically, we found a differential masking effect such that speech categorization was poorer under informational masking compared to energetic masking, despite equivalent SNRs and spectral content (*p* = 0.0226). The greater performance detriment under informational loads suggests that the warping of the acoustic-phonetic phoneme space is influenced by high-order lexical interference (Akeroyd, 2008; Myers and Blumstein, 2008).

**Figure 2:**
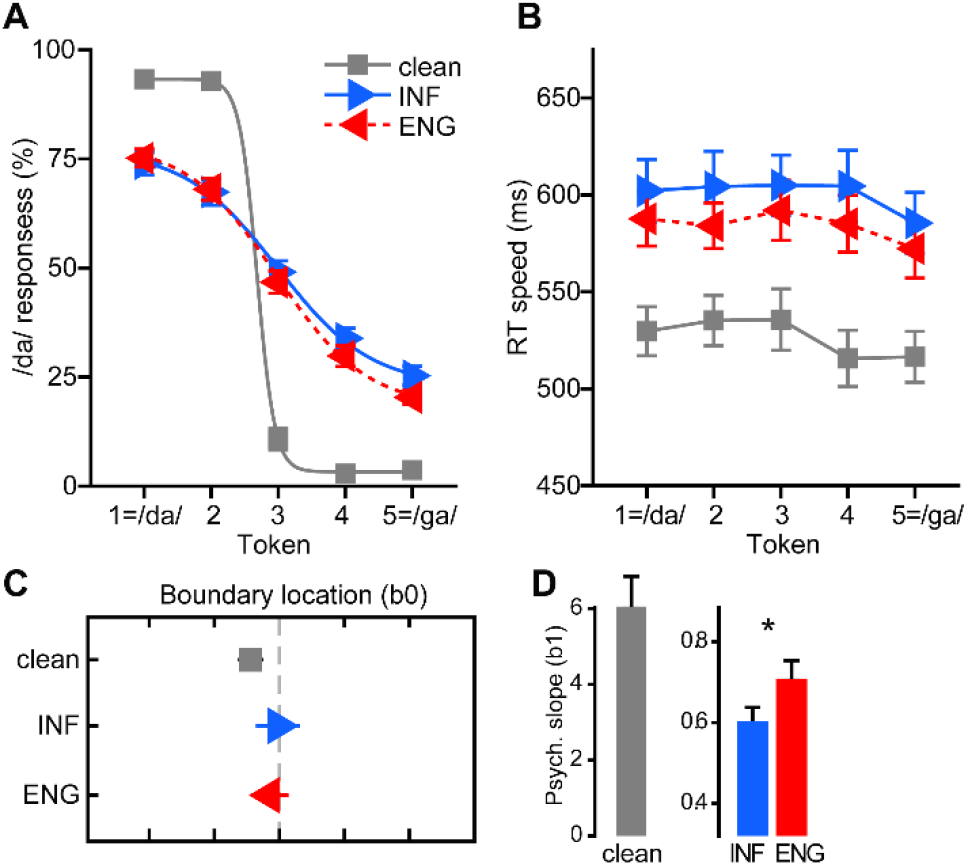
Behavioral speech categorization is differentially hindered by informational (INF) vs. energetic (ENG) acoustic interference. **(A)** Perceptual psychometric functions for phoneme identification amidst various masking noise. The clean curve shows an abrupt shift in category responses indicative of strong CP. Masking linearizes identification slopes indicative of more continuous perception but this effect various with masker type; CP is weaker under informational vs. energetic interference. **(B)** Reaction times for speech identification. Listeners are slower at labeling speech in noise overall but are faster under energetic compared to informational masking. **(C)** Location of the perceptual boundary varies little with masker type. **(D)** Psychometric slopes are shallower for informational vs. energetic masking indicating speech categorization is hindered by concurrent linguistic competition (INF masker) above and beyond acoustic interference alone (ENG masker). Error bars here and throughout = ±1 s.e.m.

Masker-related changes to RTs are shown in **Figure 2B**. RTs were modulated by masker type (*F*_2,196_ = 109.02, *p* < 0.0001) and token (*F*_4,196_ = 109.02, *p* = 0.0425). Pairwise contrasts revealed RTs for clean speech were faster than in energetic (*p* < 0.0001) and informational masking (*p* < 0.0001). Thus, noise slowed responses speeds overall. More critically, we found RTs during energetic masking were faster than during informational masking (*p* = 0.0077). The token effect was attributable to listeners’ faster decisions when labeling endpoint (Tk5) compared to categorical boundary (Tk3) tokens (*p* = 0.0361). These results are consistent with a slowing of decision speed for category ambiguous speech sounds (Bidelman and Walker, 2017; Bidelman et al., 2020a; Pisoni and Tash, 1974; Reetzke et al., 2018). The graded effect between maskers again indicates accessing perceptual categories was easier during energetic than informational noise.

Changes in the categorical boundary location are shown in **Figure 2C**. In general, listeners’ perception expectedly flipped near the midpoint of the continuum (~Tk 3) regardless of masker type. Nevertheless, an ANOVA revealed masker type moved the boundary location (*F*_2,28_ = 3.95, *p* = 0.0308). We found a small but measurable leftward shift in the categorical boundary location between clean and informational masking (*p* = 0.0104). However, informational and energetic masking did not differ (*p* = 0.6722). These findings suggest that while the overall addition of noise shifts listener’s perceptual boundary, the inherent content of the masker itself does not further bias listeners’ response. Collectively, our behavioral data suggest informational noise influences CP not via biasing listeners’ responses, *per se,* but by differentially warping and slowing access to category representations.

### 2.2 Electrophysiological data

Scalp topographies and time-domain waveforms of the cortical ERPs (i.e., channel data) are shown in **Figure 3**. Visual inspection of scalp waveforms revealed stark differences between responses in quiet and noise—expected masking effects—but no prominent effects between informational and energetic masking conditions at the scalp. Therefore, subsequent analyses were conducted on source-level data which provided a more sensitive description of neural effects.

**Figure 3:**
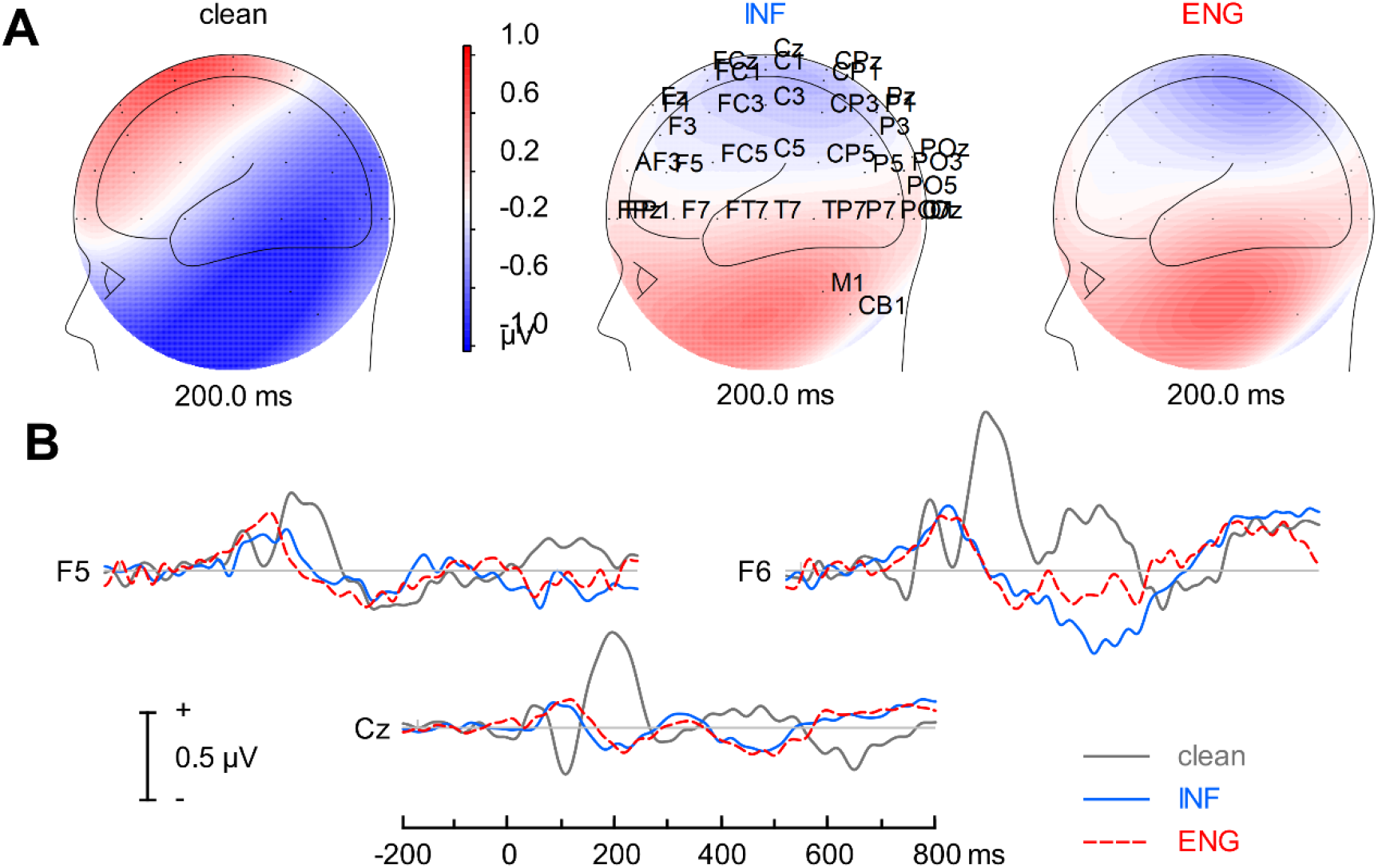
Cortical speech-ERPs. Channel-level **(A)** scalp topographies and **(B)** time-domain waveforms across noise conditions. Responses represent time-locked activity to the continuum’s speech stimuli (pooled across tokens) to illustrate noise effects. Noise reduces ERP amplitudes and prolongs latencies (e.g., Cz) with more apparent changes in morphology in frontal electrodes, particularly over right hemisphere (e.g., F6).

Cluster-based permutation testing (Maris and Oostenveld, 2007) conducted on source (AC, IFG) waveforms revealed distinct segments of neural activity that distinguished category-level information in early (~200ms) and late (400-500 ms) time windows (**Fig. 4**). For clean speech, only one early segment (172-232 ms) in right IFG differentiated ambiguous vs. prototypical tokens (Tk 3 > Tk1/5, *p*=0.0190; **Fig. 4A**). More extensive differences in categorical coding were observed during energetic and informational masking, with additional clusters observed in multiple ROIs and time segments across the epoch window (**Figs. 4B,C**). In the presence of energetic interference—perceptually tasking but devoid of linguistic content—token-wise differences were observed first in early (282–328 ms) left IFG (Tk1/5 >Tk3, *p*=0.0250) followed later by left AC (Tk1/5 >Tk3, 540-578 ms; *p*=0.030) (**Fig. 4B**). During informational masking, speech categories were distinguishable during an early period in bilateral IFG and left AC, with additional clusters emerging later in time within left AC and right IFG (Tk 3 >Tk 1/5, all *p*’s ≤ 0.055) (**Fig. 4C**).

**Figure 4:**
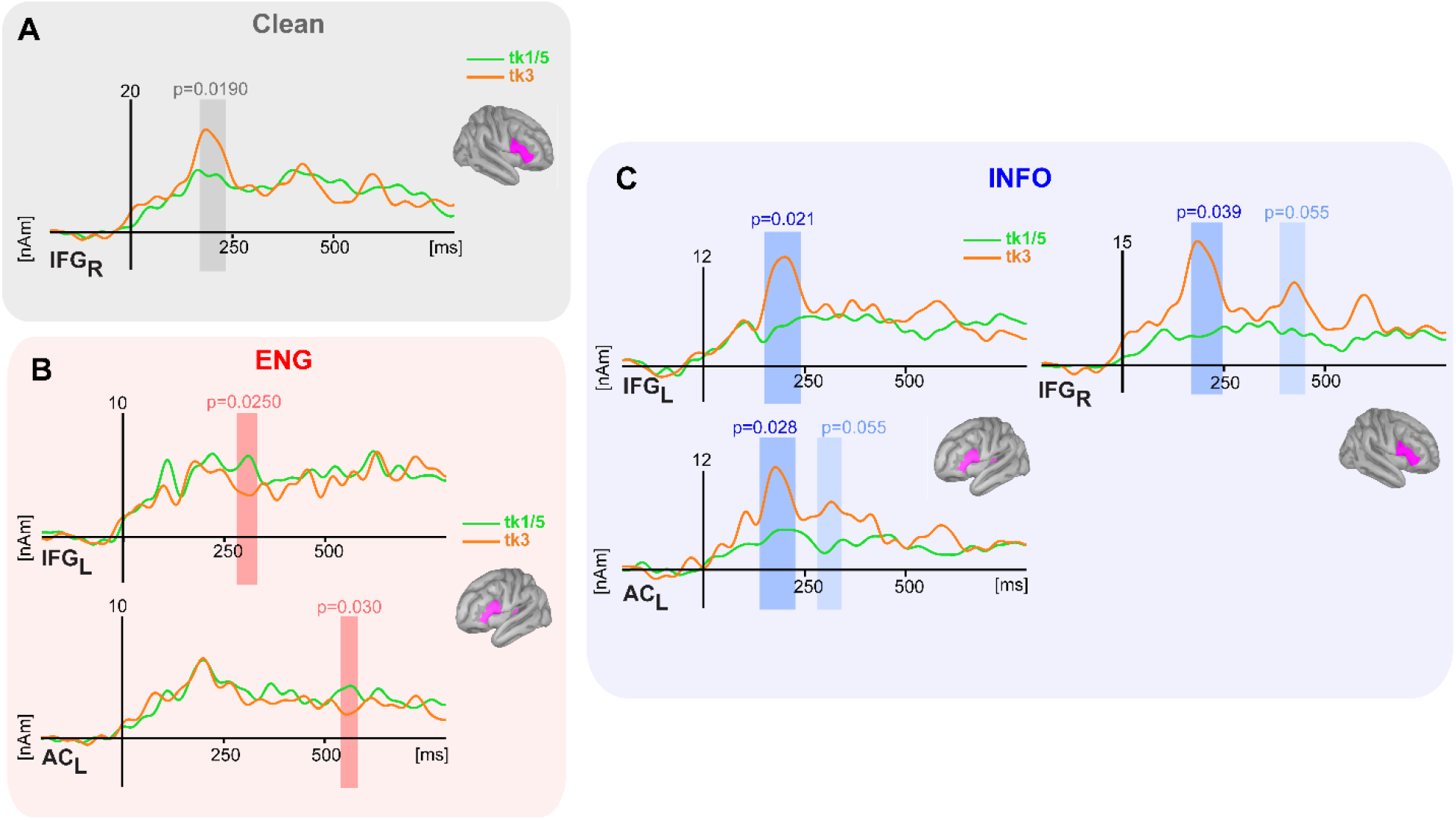
Cluster-based permutation results jointly comparing the strength of categorical processing between prototype (green) and boundary (orange) tokens across masker types and ROIs. Shading = segments where ERPs show category-level coding (i.e., Tk15 ≠ Tk3, *p*<0.05, corrected). (**A**) For clean speech, early right-lateralized IFG activity shows categorical organization. (**B**) During energetic masking, categorical speech processing emerges during both early and late stages in AC and IFG. (**C**) During informational masking, speech categories are distinguishable in early bilateral IFG and left AC, with additional later clusters in left AC and right IFG.

**Figure 5** shows grand mean ERP source waveforms in bilateral AC and IFG. These responses were pooled across tokens to assess masker-specific effects. A preliminary running *t-*test (Guthrie and Buchwald, 1991) identified a time segment between ~200-500 ms that differentiated responses in information vs. energetic masking. AC effects tended to occur in an early time window (175-225), whereas IFG showed unique masker-related effects in a later period (350-400 ms). As expected from prior speech-ERP studies (Bidelman and Howell, 2016; Billings et al., 2009), the introduction of noise decreased response amplitudes relative to clean speech in all brain regions. However, informational masking further reduced amplitudes compared to energetic masking in left AC and bilateral IFG. An ANOVA showed source amplitudes varied with masker type (*F*_1,98_ = 9.71, *p* = 0.0024) but not ROI (*F*_3,98_ = 1.10, *p* = 0.3510). The masker effect was attributable to lower amplitudes under informational compared to energetic masking (*p* < 0.0001). The lack of ROI effect (and masker x ROI interaction) suggests informational masking weakened speech processing above and beyond its energetic counterpart in both auditory and linguistic cortex.

**Figure 5:**
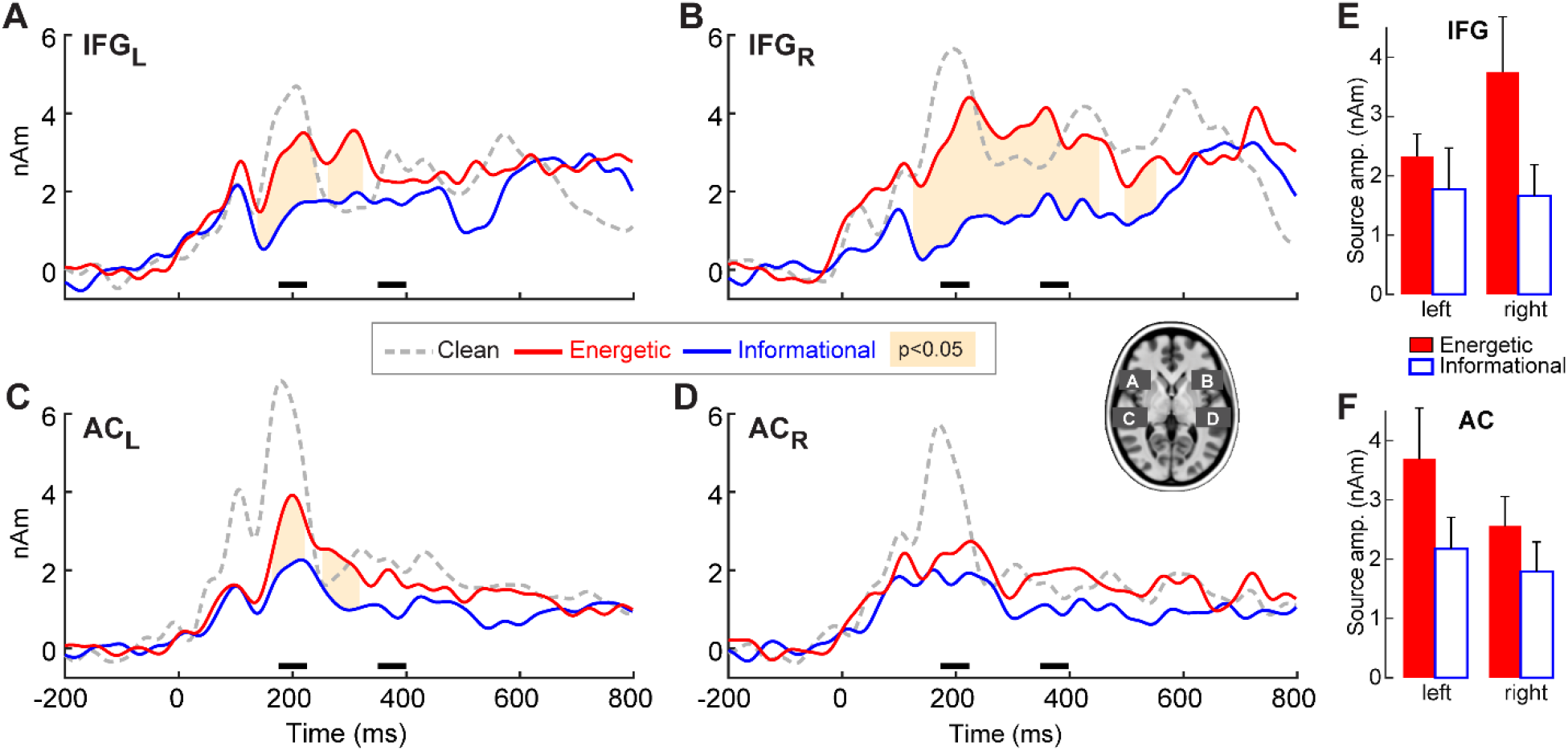
Speech-ERP source waveforms as a function of masker extracted from bilateral auditory cortex (AC) and inferior frontal gyrus (IFG). **(A)** Left IFG, **(B)** Right IFG, **(C)** Left AC, and **(D)** Right AC. Shaded regions denote time segments where responses differ between masker type (running *t-*test, *p*<0.05) (Guthrie and Buchwald, 1991). (**E-F**) Source waveform amplitudes between the informational and energetic masking conditions in IFG and AC. AC and IFG amplitudes were extracted from and early (~200 ms) and late (~400 ms) time windows, respectively (see bars along abscissa, panels A-D). Neural responses are weaker when categorizing speech during informational vs. energetic masking indicating linguistic interference hinders the acoustic-phonetic conversion process of CP.

### 2.3 Functional connectivity

We quantified the strength of frequency-specific connectivity between brain regions using Granger causality (**Fig. 6**). An ANOVA (*F*-test) conducted on the time-frequency connectivity maps between all pairwise ROIs revealed a significant cluster in the AC_left_ → IFG_right_ connection (*p* = 0.004) in the low gamma band (35-40 Hz) between 100-300 ms (**Fig. 6A)**. That is, the strength of cross hemispheric temporo-frontal connectivity differed across masking conditions. Post hoc permutation tests showed this effect was attributed to stronger gamma-band connectivity in clean relative to energetic masking (*p* = 0.001); AC_left_→ IFG_right_ neural signaling did not differ between the informational and energetic noise conditions (*p* = 0.3843; **Fig. 6B-C**). These results suggest right IFG (and information broadcast from left AC) plays an active role in categorizing degraded speech, a circuit which is further modulated with task difficulty.

**Figure 6:**
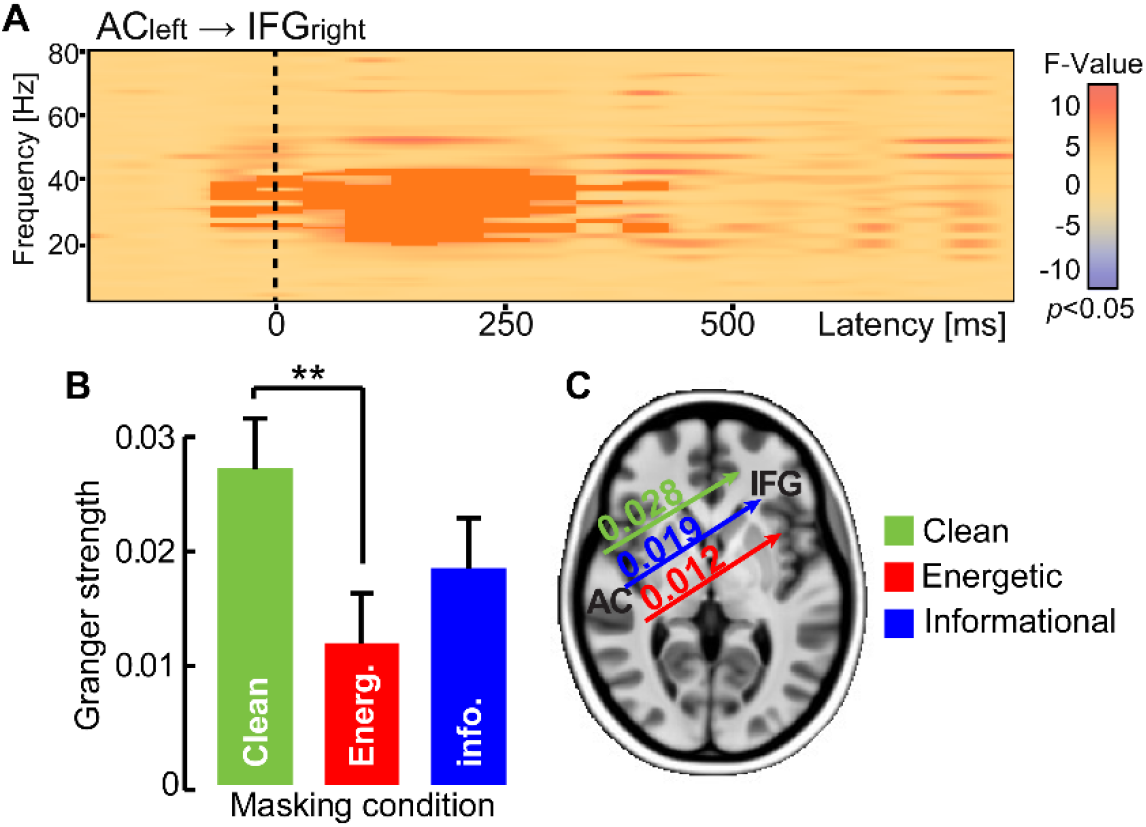
Functional connectivity (Granger causality) reveals directed neural signaling from left AC to right IFG differentiates speech categorization under different forms of noise. (**A**) The spectrographic map shows frequency-specific AC_left_→IFG_right_ connectivity across frequency and time of the epoch window. Dark shading, clusters where connectivity strength differed between noise conditions (*F*-test; *p*<0.05). (**B-C**) Connectivity strength in the low gamma band (35-40 Hz; 100-300 ms) is higher when classifying clean speech and weakens during energetic masking. Connectivity is intermediate for informational masking. **p< 0.01

### 2.4 Brain-behavior relationships

We ran Pearson’s correlations to assess relations between brain responses (each ROI) and behavior. The correlations compared the difference in source amplitudes between informational and energetic masking (e.g., ∆IFG = IFG_I_ – IFG_E_) with the homologous difference in behavioral reaction times (∆RT= RT_I_ - RT_E_). Thus, positive ∆ amplitudes indicate larger neural responses (in a given ROI) for informational vs. energetic masking (which rarely occurred, cf. **Fig. 5**); likewise, a positive ∆RT indicates slower decision speeds during informational masking. We found a negative correlation between differential responses in left IFG (∆IFG_left_) and behavioral RTs (r = −0.63, p = 0.012; **Fig. 7**), suggesting poorer IFG encoding of speech phonemes under informational loads was related to slower categorization decisions. No other ROIs showed correlations with behavioral RTs or psychometric slopes (data not shown).

**Figure 7:**
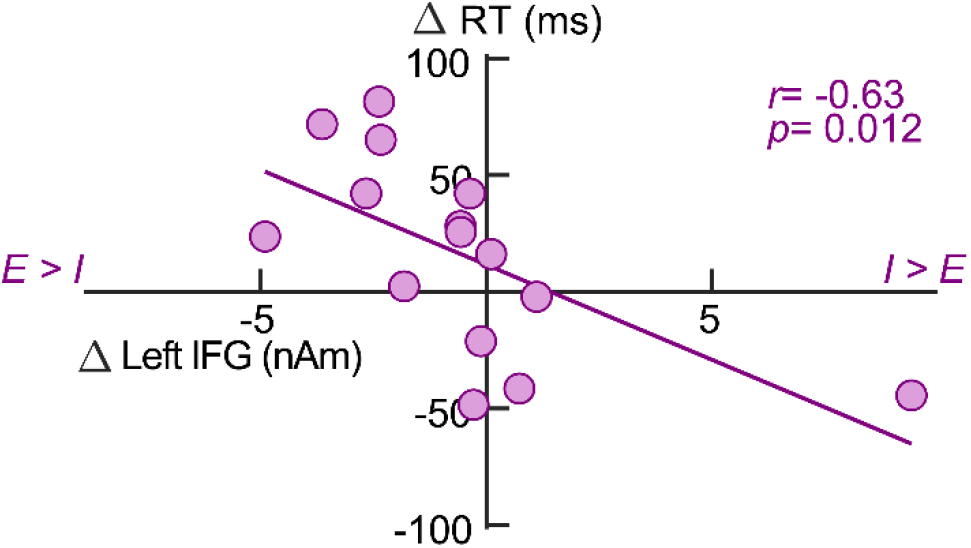
Pearson’s correlation showing the relationship between masker-related changes in left inferior frontal gyrus and behavioral reaction times. ∆IFG reflects the change in source amplitude within left IFG between informational and energetic masking (i.e., ∆IFG = IFG_I_ – IFG_E_). Similarly, ∆RT reflects the change in behavior (i.e., ∆RT = RT_I_ - RT_E_). Positive ∆IFG and ∆RT indicate larger neural responses and slower decision speeds during informational relative to energetic loads. The negative association indicates listeners who were more susceptible to lexical influences at the neural level also show less benefit between I and E behaviorally. I= informational masking; E=energetic masking.

## 3. DISCUSSION

By measuring behavior and brain activity (ERPs, functional connectivity) during speech categorization in different acoustic noises (i.e., spectrally matched energetic vs. informational masking), our data reveal three primary findings: (1) behaviorally, informational masking weakens speech phoneme identification above and beyond acoustic/auditory masking; (2) low-level auditory cortical activity not only codes speech categories but is susceptible to higher-order lexical interference; (3) identifying speech amidst noise recruits a cross hemispheric circuit (AC_left_ → IFG_right_) whose engagement varies according to task difficulty. Taken together, our results provide corroborating evidence for top-down influences on the early acoustic-phonetic analysis of speech.

Behaviorally, we found listeners’ speech identification was modulated by the specific type of noise interference. Psychometric slopes were steeper when categorizing clean relative to noise-degraded speech, consistent with well-known masking effects on CP that emerge at lower (i.e., negative) SNRs (Bidelman et al., 2019; Bidelman et al., 2020a). More critically however, we found that informational masking yielded shallower CP slopes than energetic masking. Our energetic masker was the informational masker reversed in time, meaning both maskers had the same spectral profile and were matched in overall SNR—only their linguistic status differed. Consequently, the weaker performance under informational loads suggests that added lexical interference hinders speech categorization above and beyond acoustic interference alone—a top-down influence on categorical processing (Bidelman et al., 2020b; Ganong, 1980; Gow et al., 2008; Myers and Blumstein, 2008; Scott and Wise, 2004). However, the fact that the categorical boundary did not shift appreciably with additional informational masking suggests lexical interference does not bias listeners in their responses, *per se*. Instead, our behavioral data suggest informational noise influences speech categorization by differentially warping and slowing access to the category representations of speech (e.g., Fig. 2B, 2D).

In this vein, RTs for categorization were longer with the addition of the noise, suggesting slower access to speech representations in challenging acoustic conditions (Bidelman et al., 2019; Bidelman et al., 2020a; Pisoni and Tash, 1974). More interesting, however, is the differential masking effect on decision speeds. RTs further slowed when categorizing speech under informational relative to energetic loads. This suggests accessing categories is further stalled when the brain is faced with juggling the lexical interference from informational masking. Listeners who were more susceptible to these top-down influences neutrally, specifically within left IFG, tended to have slower RTs (Fig. 7). This is consistent with previous work showing individuals who are more susceptible to lexical influences experience greater processing difficulties parsing concurrent speech (e.g., as in our informational masking condition) (Lam et al., 2017; Price et al., 2019).

Informational masking effects on behavioral CP could manifest via bottom-up (through direct influences on perception of the signal) or top-down processes (through modulation of post-perceptual processing from higher order brain areas) (McClelland and Elman, 1986; Norris et al., 2000). Our neuroimaging data help resolve the level at which lexical effects from informational masking occurs. Electrophysiologic results demonstrated that informational masking had a more detrimental impact to categorical processing across both auditory (AC) and linguistic (IFG) brain areas. Informational effects were particularly evident in bilateral inferior frontal gyrus, consistent with the role of IFG in speech identification, particularly under degraded conditions (Bidelman and Walker, 2019; Du et al., 2014; Myers et al., 2009). More critically however, lexical masking was observed in (left) auditory cortex. These data suggest that early auditory sensory areas and their acoustic-phonetic representations for speech are directly susceptible to lexical influence, conventionally considered a higher-level construct in speech perception (Myers et al., 2009). This notion is further supported by our correlational analysis which showed that informational-related changes in left AC responses predicted changes in behavioral RTs.

The link between left IFG and behavior supports previous findings suggesting IFG drives listeners’ RTs in speech categorization tasks (Binder et al., 2004). This suggests a link between left IFG activation and lexical responses in auditory tasks. As our RTs are indicative of categorical processing, they signify slower access to categories with neural correlates in IFG, instead of responses related to SNR differences. This further demonstrates activation of a linguistic network to disambiguate speech from noise (Bidelman and Howell, 2016; Du et al., 2014).

Time course analysis revealed stronger neural responses to category vs. ambiguous speech tokens in purely energetic, non-linguistic noise (Fig. 4B). This is consistent with the notion that phonetic features (e.g., speech categories) are more resilient to acoustic interference than their non-phonetic acoustic counterparts (Bidelman et al., 2019; Bidelman et al., 2020a; Lewis and Bidelman, 2020). The leftward bias and temporal dynamics (IFG preceding AC) of this effect further imply that while categorical representations are evident in both higher-order (linguistic) and lower-level (auditory) cortex (e.g., Bidelman and Lee, 2015; Binder et al., 2004; Chang et al., 2010; Feng et al., 2018; Mankel et al., 2020; Myers et al., 2009), categories might be first decoded in IFG prior to their emergence in AC. Along these lines, IFG leading AC activation has been previous observed in degraded speech perception tasks (Bidelman and Dexter, 2015) and is consistent with notions that higher order speech centers exert an inhibitory influence on concurrent auditory representations to prevent interference from nonlinguistic cues (Dehaene-Lambertz et al., 2005; Liberman et al., 1981). Such preemptive, top-down influences would be beneficial to identify speech in noise. Although, we note that top-down (i.e., IFG→AC) connectivity did not differentiate neural responses under different masking conditions so this interpretation remains speculative.

We found additional early (~250 ms) and late (~450 ms) engagement of right IFG during categorical processing amidst informational masking. This suggests that in addition to a left lateralized auditory-linguistic circuit (i.e., IFG↔AC), the brain recruits a widely distributed and bilateral network to decode target speech categories under the added constraints of concurrent lexical interference. This notion is further bolstered by the cross-hemisphere transmission and modulation of AC_left_→IFG_right_ connectivity with the differing demands across noise stimuli. Previous EEG studies have similarly shown increased right hemisphere engagement for noise-degraded speech, particularly areas in frontal lobe (Bidelman and Howell, 2016; Mahmud et al., 2020; Price et al., 2019). The current data are at broadly consistent with these previous findings.

Right IFG has previously been shown to play a role in inhibitory control during response selection (Garavan et al., 1999). Meanwhile, others have shown right IFG activation relates with attentional control and detection of important task-relevant cues (Hampshire et al., 2009; Hampshire et al., 2010). Further research has demonstrated right IFG plays a role in processing phonology and semantics (Hartwigsen et al., 2010). Neuroimaging work has also demonstrated increased engagement of right hemisphere PAC and IFG with decreasing speech SNR (Bidelman and Howell, 2016; Doeller et al., 2003). This could be due to the signal becoming so degraded during the energetic noise that inhibition becomes unnecessary, as the noise has already warped the percept to a degree that it no longer matches the internalized category template of the phoneme. Alternatively, masking could increase cognitive load causing listeners to partially disengage with the task and direct attention elsewhere (e.g., to the noise rather than target phoneme). Thus, the observed upregulation of right IFG could relate to the increased attentional demands of classifying noise-degraded speech. Under this account, we may have expected even larger right IFG activation under informational masking, an arguably more challenging listening condition; however, this was not what we observed (**Fig. 5**). Alternatively, increased fronto-temporal interactions might increase in situations of degraded sensory input as part of predictive coding or compensatory strategy (Cope et al., 2017; Price et al., 2019). The fact that right IFG differentiated prototypical and boundary stimuli in the clean and informational masking conditions provides additional evidence that right IFG may provide predictive coding, even if it might fail in some conditions. Still, we might then have expected more top-down (IFG→AC) connectivity which is not what we observed. Perhaps due to the relative ease of processing speech in quiet, this connection, and therefore predictive detection of task-related cues, is strengthened in that condition relative to noise. The informational condition also has two streams of linguistic information (target phoneme and masker words). Increased IFG activation could be related to trying to predict both streams, causing greater connectivity from left AC compared to the energetic masking condition. Future studies are needed to clarify the function of right IFG and its role in the auditory-linguistic speech network.

Broadly, the differential effect of informational masking here supports other observations in the literature, namely, that categorization of speech occurs in higher-order and lower-order brain regions, and converging evidence of lexical effects on categorical processing (Bidelman et al., 2020b; Ganong, 1980; Myers and Blumstein, 2008). Lexical effects, while perhaps governed by higher-order brain areas, begins influencing the neural encoding of the signal as early as auditory cortex. Together, these findings suggest tight coupling in the fronto-temporal brain areas associated with speech perception allow for categorization to occur even when there is competing noise.

## 4. MATERIALS & METHODS

### 4.1 Participants

Fifteen young adults (age 24.8 ± 5.0 years; 8 females) participated in this experiment. All had normal audiometric thresholds (thresholds ≤20 dB HL; 250-8000 Hz) and were native speakers of English; we selected native listeners as speech in noise detriments occur in bilinguals in their non-native language (Bidelman and Dexter, 2015; Krizman et al., 2017; Lucks Mendel and Widner, 2016; Tabri et al., 2015). Listeners were largely right-handed (mean 73% laterality; Edinburgh Handedness Inventory: Oldfield, 1971) and had obtained some level of college education (17.4 ± 3.5 years). Music training improves speech-in-noise perception (Bidelman and Yoo, 2020; Parbery-Clark et al., 2009; Yoo and Bidelman, 2019). Thus, all participants were required to have minimal (≤3 years) formal music training throughout their lifetime. Each gave written informed consent in compliance with a protocol approved by the IRB at the University of Memphis.

### 4.2 Stimuli & task

We used a synthetic five-step CV syllable continuum spanning from /da/ to /ga/ (based on placed of articulation). Stimulus morphing was achieved by altering the F2 formant region in a stepwise fashion using the STRAIGHT software package (Kawahara et al., 2008) based on original speech materials described in Nath and Beauchamp (2012) (https://openwetware.org/wiki/Beauchamp:Stimuli). Each stimulus was 500 ms in duration but was truncated to 350 ms for task presentation (**Figure 1**).

In addition to a “clean” (i.e., no noise) condition, phoneme identification was measured in two forms of acoustic noise designed to create more (informational masker) or less (energetic masker) linguistic interference. The informational masker was a two-talker babble (Niemczak and Vander Werff, 2019); that same masker was reversed in time to remove the linguistic components and create a purely energetic masker. The babble consisted of two female voices reading aloud from printed materials, with the long-term average speech spectra of the maskers equalized in 1/3 octave bands (±5 dB). The energetic masker was created by time-reversing the babble in Audacity (v 2.2.3; www.audacityteam.org/), thus matching its long-term spectral profile. Masking conditions were presented in separate blocks and pseudo-randomized using a Latin Square counterbalance (Bradley, 1958).

We delivered speech stimuli binaurally through ER-2 insert earphones (Etymotic Research, Elk Grove Village, IL) at 74 dBA SPL in quiet and the presence of energetic and informational maskers. Maskers were presented at 76 dBA SPL, for a signal to noise ratio (SNR) of −2 dB. We selected this SNR based on previous findings that speech categorization is resilient to noise down to ~0 dB SNR (Bidelman et al., 2020a) and extensive pilot testing, which confirmed −2 dB SNR was adequate to decrease the steepness of listeners’ psychometric function. Stimuli were calibrated using a Larson Davis sound pressure level meter (model LxT + mic 377B20). Sound delivery was controlled by a custom MATLAB program coupled to a TDT RP2 digital signal processor (Tucker-Davis Technologies, Alachua, FL).

During the behavioral task, listeners heard 150 trials of each CV stimulus per noise block (total = 750 per block). Listeners rapidly responded to phonemes on each trial with a binary response on the keyboard (“da” or “ga”). We instructed them to respond as quickly and accurately as possible. Following their behavioral response, the interstimulus interval was jittered randomly between 800 and 1000 ms (20 ms steps, uniform distribution). We allowed breaks between blocks to avoid listening fatigue. Percent identification and reaction times (RTs) were logged.

### 4.3 EEG recording procedures

Continuous EEGs were recorded during the active speech identification task from 64 sintered Ag/AgCl electrodes at standard 10-10 scalp locations (Neuroscan QuikCap array) (Oostenveld and Praamstra, 2001). Continuous data were digitized using a sampling rate of 500 Hz (SynAmps RT amplifiers; Compumedics Neuroscan) and an online passband of DC-200 Hz. Electrodes placed on the outer canthi of the eyes and the superior and inferior orbit monitored ocular movements. Contact impedances were maintained <10 kΩ during data collection. During acquisition, electrodes were referenced to an additional sensor placed ~1 cm posterior to the Cz channel. Data were re-referenced to the common average for analysis.

EEG pre-processing was performed in BESA® Research (v7) (BESA, GmbH). Ocular artifacts (saccades and blinks) were first corrected in the continuous EEG using a principal component analysis (PCA) (Picton et al., 2000). EEGs were then bandpass filtered from 1-20 Hz (−48 dB/Hz roll-off), baselined, epoched from −200 − 800 ms, and averaged per token to derive ERPs for each stimulus condition.

To reduce the dimensionality of the data and permit functional connectivity analysis between brain regions of interest (ROIs), we transformed the scalp potentials into source space using virtual source montaging in BESA (Scherg et al., 2002). This process applied an optimized spatial filter to all electrodes that calculated their weighted contribution to the scalp recorded ERPs. This provided an estimate of activity at various source locations within the head while reducing overlapping activity from other brain regions (for details, see Scherg and Ebersole, 1994; Scherg et al., 2002). Based on the Talairach atlas (Talairach and Tournoux, 1988), we seeded regional sources in bilateral Auditory Cortex (AC) [*x,y,z* (mm); *left:* −50.4, − 17.3, 11.6; *right*: 50.4, −17.4, 12.5] and Inferior Frontal Gyrus (IFG) [*left*: −33.8, 26.5, 2.1; *right:* 33.8, 26.5, 2.1]. Regional sources consist of three dipoles describing current flow (units nAm) in the X, Y, and Z planes. We combined the three orientations at each location (i.e., L2-norm) to compute the overall power generated in each ROI. Focusing on source waveforms allowed us to reduce each listener’s 64 electrode data to four source channels describing neural activity in the auditory-linguistic pathways (i.e., bilateral IFG and AC).

### 4.4 Behavioral data analysis

Identification scores were fit to a sigmoidal function *P* = 1/[1 + e^−*β1*(*x*−*β0*)^], where *P* is the proportion of trials identified as a given CV syllable, *x* is the step number along the stimulus continuum, and *β0* and *β1* the location and slope of the logistic fit estimated using non-linear least-squares regression. Comparisons of *β0* location between noise conditions reveals changes in the perceptual boundary; whereas changes in *β1* demonstrate noise-related changes in the strength of speech categorization, with steeper psychometric functions (i.e. larger *β1*) indicating stronger categorical hearing.

Behavioral reaction times (RTs) were computed as the median response latency for each token in each condition. RTs outside of 250-2000 ms were considered outliers (i.e., guesses or lapse of attention) and were excluded from RT analysis.

### 4.5 Electrophysiological data analysis

#### 4.5.1 Cluster-based permutation statistics on source waveforms

To first evaluate whether source ERPs showed categorical organization, we averaged response amplitudes to tokens at the endpoints of the continuum and compared this combination to the ambiguous midpoint token (e.g., Bidelman and Walker, 2017; Liebenthal et al., 2010). This contrast [i.e., mean (Tk1, Tk5) vs. Tk3] minimizes stimulus-related differences in the ERPs and isolates categorical/perceptual processing. We then used cluster-based permutation statistics (Maris and Oostenveld, 2007) to test for categorical effects (i.e., Tk1/5 vs. Tk3) in the source ERPs. A two-stage analysis was conducted in BESA Statistics 2.0. First, we computed a series of paired *t*-tests contrasting source amplitudes at every time point in the epoch window (−200 – 800 ms). This preliminary step identified clusters both in time (adjacent samples) and space (sources) where responses differed between conditions (*p*<0.05). The initial clustering based on the preliminary parametric t-tests was then followed by permutation testing on these clusters. Cluster values were based on the sum of all *t*-values of data points within a given cluster. Significant differences were determined by generating surrogate clusters from *N*=1000 resamples of the data and permuting between stimulus conditions (e.g., Oostenveld et al., 2011). This identified contiguous time samples where the stimulus conditions were not interchangeable (i.e., Tk1/5 ≠ Tk 3; *p* < 0.05). Importantly, the clustering process ensured results are corrected for multiple comparisons across the aggregate of all time points and source channels by controlling the familywise error rate.

#### 4.5.2 ERP quantification of noise-related effects

The initial cluster analysis revealed early and late time windows coding category-level information (see Fig. 4). In subsequent quantification of noise-related effects, we measured amplitudes from the time-domain source waveforms associated with early AC (175-225 ms) vs. IFG (350-400 ms) activity, respectively. For these analyses, we pooled responses across tokens of the speech continuum to focus on noise-related changes in speech processing. Average wave amplitude was computed from each time segment for each ROI and masker condition.

#### 4.5.3 Functional connectivity (Granger causality)

To evaluate functional connectivity between brain regions, we measured directed information flow between bilateral AC and IFG using Granger Causality (GC) (Geweke, 1982; Granger, 1969). We used BESA Connectivity (v1) to compute GC in the frequency domain (Geweke, 1982) using nonparametric spectral factorization on the single-trial time-frequency maps (Dhamala et al., 2008). The frequency decomposition was based on complex demodulation (Papp and Ktonas, 1977). From cleaned EEGs, single trial data were recomputed using full band (1-80 Hz, with 60 Hz notch filter) responses, pooling across the tokens to observe the effects of masking on connectivity. We imposed an additional layer of artifact thresholding such that >85% of trials per participant remain in the analysis. Signal *X* is said to “Granger-cause” signal *Y* if past values of *X* contain information that predict *Y* above and beyond information contained in past values *Y* alone. Importantly, GC can be computed directionally (e.g., *X→Y* vs. *Y→X*) to infer causal flow between interacting brain signals. Connectivity was computed in both the bottom-up (AC→IFG) and top-down (IFG→PAC) directions as well as the ipsilateral and contralateral (crossed-hemisphere) connections (e.g., AC_left_→IFG_right_). These analyses provided a spectrogram-like representation of GC strength between pairwise sources across the entire epoch window and frequency bandwidth of the EEG. We then conducted a one-way ANOVA on connectivity data in BESA Statistics (v2.0). This identified specific frequency channels and directional connections where Granger strength was modulated by the type of noise interference (i.e., clean vs. informational vs. energetic masking).

### 4.6 Statistical analysis

Unless otherwise noted, dependent measures were analyzed using a two-way mixed model ANOVA (subjects = random factor) with fixed effects of ROI (4 levels: left/right AC and IFG) and masker type (3 levels: clean, informational, energetic) (PROC GLIMMIX, SAS® 9.4; SAS Institute, Inc.). Tukey-Kramer adjustments controlled for Type I error inflation for multiple comparisons. The α-level for significance was p = 0.05. We used Pearson’s correlations to assess associations between EEG and behavioral responses.

## Acknowledgements

Work supported by the National Institute on Deafness and Other Communication Disorders of the National Institutes of Health under award number R01DC016267 (G.M.B.). We thank Kelsey Mankel for comments on earlier versions of this manuscript. Requests for data and materials should be directed to G.M.B.

